# Altered temporal sequence of transcriptional regulators in the generation of human cerebellar granule cells

**DOI:** 10.1101/2021.01.17.427030

**Authors:** Hourinaz Behesti, Arif Kocabas, David E. Buchholz, Thomas S. Carroll, Mary E. Hatten

## Abstract

Brain development is regulated by conserved transcriptional programs across species, but little is known about divergent mechanisms that create species-specific characteristics. Among brain regions, the cerebellum is now recognized to contribute to human cognitive evolution having a broad range of non-motor cognitive functions in addition to motor control. Emerging studies highlight the complexity of human cerebellar histogenesis, compared with non-human primates and rodents, making it important to develop methods to generate human cerebellar neurons that closely resemble those in the developing human cerebellum. Here we report a rapid and simple protocol for the directed derivation of the human ATOH1 lineage, the precursor of excitatory cerebellar neurons, from human pluripotent stem cells (hPSC), and strategies to decrease culture variability; a common limitation in hPSC studies. Upon transplantation into juvenile mice, early postmitotic hPSC-derived cerebellar granule cells migrated along glial fibers and integrated into the cerebellar cortex. By Translational Ribosome Affinity Purification (TRAP)-seq, the ATOH1 lineage most closely resembled human cerebellar tissue in the second trimester. Unexpectedly, TRAP-seq identified a heterochronic shift in the expression of RBFOX3 (NeuN) and NEUROD1, which are classically associated with differentiated neurons, within granule cell progenitors (GCPs) in the human external granule layer. This molecular divergence may provide the mechanism by which the GCP pool persists into year two post birth in humans, but only lasts for two weeks in mice. Our approach provides a scalable *in vitro* model of the human ATOH1 lineage that yields cerebellar granule cells within 48 days as well as a strategy for identifying uniquely human cellular and molecular characteristics.

## Introduction

Understanding the development of the human brain is an emerging area of neuroscience. The human cerebellum is now recognized to contribute to cognitive functions (Allen et al., 1997; Carta et al., 2019; Fiez, 1996; Schmahmann et al., 2019; Stoodley et al., 2017; Wagner et al., 2017), in addition to a critical role in motor control and motor learning. As one of the most ancient cortical regions, the cerebellum also appears central to human cognitive evolution, having rapidly expanded in absolute size, and relative to the neocortex in apes and humans (Barton and Venditti, 2014). Recent studies provide evidence of the complexity of human cerebellar histogenesis, compared with non-human primates and rodents (Haldipur et al., 2019), making it important to develop methods to generate human cerebellar neurons to model human development and disease. The cerebellar cortex develops from rhombomere 1 (Wingate and Hatten, 1999) with a primary germinal zone that produces the Purkinje neuron and interneurons, and a secondary germinal zone, marked by the *ATOH1* transcription factor, that emerges from the rhombic lip and generates cerebellar granule cells (GCs) (Hatten and Heintz, 1995). Importantly, prior to the specification of granule cell progenitors (GCPs), the ATOH1 lineage gives rise to a subset of hindbrain nuclei and cerebellar nuclei in mice (Machold and Fishell, 2005; Wang et al., 2005). While previous studies have mostly focused on generating human Purkinje cells (Buchholz et al., 2020; Erceg et al., 2010; Muguruma et al., 2015; Nayler et al., 2017; Silva et al., 2020; Wang et al., 2015; Watson et al., 2018) from human pluripotent stem cells (hPSCs), little attention has been given to defining the molecular pathways that generate human GCs and other rhombic lip derivatives. The importance of understanding the human ATOH1 lineage is underscored by the fact that GCPs are a known cell of origin for medulloblastoma; the most common metastatic childhood brain tumor (Marino et al., 2000), and GCs are implicated in neurodevelopmental disorders including autism (Bauman, 1991; Kloth et al., 2015; Menashe et al., 2013).

Here we report a rapid and simple protocol for the directed derivation of the human ATOH1 lineage, the precursor of excitatory cerebellar neurons, by the sequential addition of six factors, in a chemically defined culture medium. Using transgenic reporter lines and Translational Ribosome Affinity Purification (TRAP)-seq adapted to hPSCs, we tracked developmental gene expression in culture in a lineage specific manner. Compared to previously reported studies, this method accelerates the production of *ATOH1^+^* cells (day in vitro (DIV)16 versus DIV 35 in previous studies (Muguruma et al., 2015), with a dramatic increase in yield (80% versus 17%). Strategies to overcome culture variability in terms of gene expression, a common limitation of hPSC-derived models, included the addition of BMP7 to stabilize *ATOH1* gene expression, patterning of cells from a single cell-stage, and growth on transwell membranes where cells have access to medium in two dimensions (above and below). The translational profile of the hPSC-derived ATOH1 lineage most closely resembled human cerebellar tissue in the second trimester compared to other brain regions. Finally, we report the discovery of a heterochronic shift in the expression transcriptional regulators (RBFOX3 (NeuN) and NEUROD1) in the progenitor zone of the human EGL. NeuN and to a large extent NeuroD1 are expressed in postmitotic neurons in vertebrates (Miyata et al., 1999; Mullen et al., 1992). This molecular divergence may provide the mechanism whereby the GCP pool persists into year two post birth in humans, but only for two weeks in mice.

## Results

### Directed derivation of the human ATOH1 lineage from hPSCs

Addition of dual-SMAD inhibitors for neuralization (Chambers et al., 2009), followed by the addition of Fibroblast Growth Factor (FGF) and a small molecule agonist of WNT signaling (CHIR99021, “CHIR” from hereon) for posteriorization, in a chemically defined serum-free medium, induced the expression of anterior hindbrain markers *EN2, MEIS2, GBX2* and repressed midbrain (*OTX2*), and spinal cord-level (*HOXA2*) markers (Supp Fig. 1A) by DIV11. FGF+CHIR treatment was superior at inducing mid/hindbrain markers (*EN2, GBX2*) and reducing mid/forebrain markers (*OTX2/PAX6*), compared to a previously reported combination of insulin+FGF2 (Muguruma et al., 2015) (Supp Fig. 1A). Empirical testing of the timing, duration, and concentrations of CHIR+FGF, necessary for a “cerebellar territory” expression profile of EN2^+^;GBX2^+^;OTX2^-^, revealed that the addition of 2.5μM CHIR from DIV1 until at least DIV9-11 (Fig. 1A, Supp. Fig. 1B) was necessary and that dual-SMAD inhibition until DIV7 was sufficient (Supp. Fig. 1B and data not shown). While both FGF8b (Chi et al., 2003; Guo et al., 2010; Martinez et al., 1999) and FGF2 (Muguruma et al., 2015) treatments induced cerebellar territory, FGF2 induced higher *EN2* levels and improved cell survival (Supp Fig. 1B).

**Figure 1.**
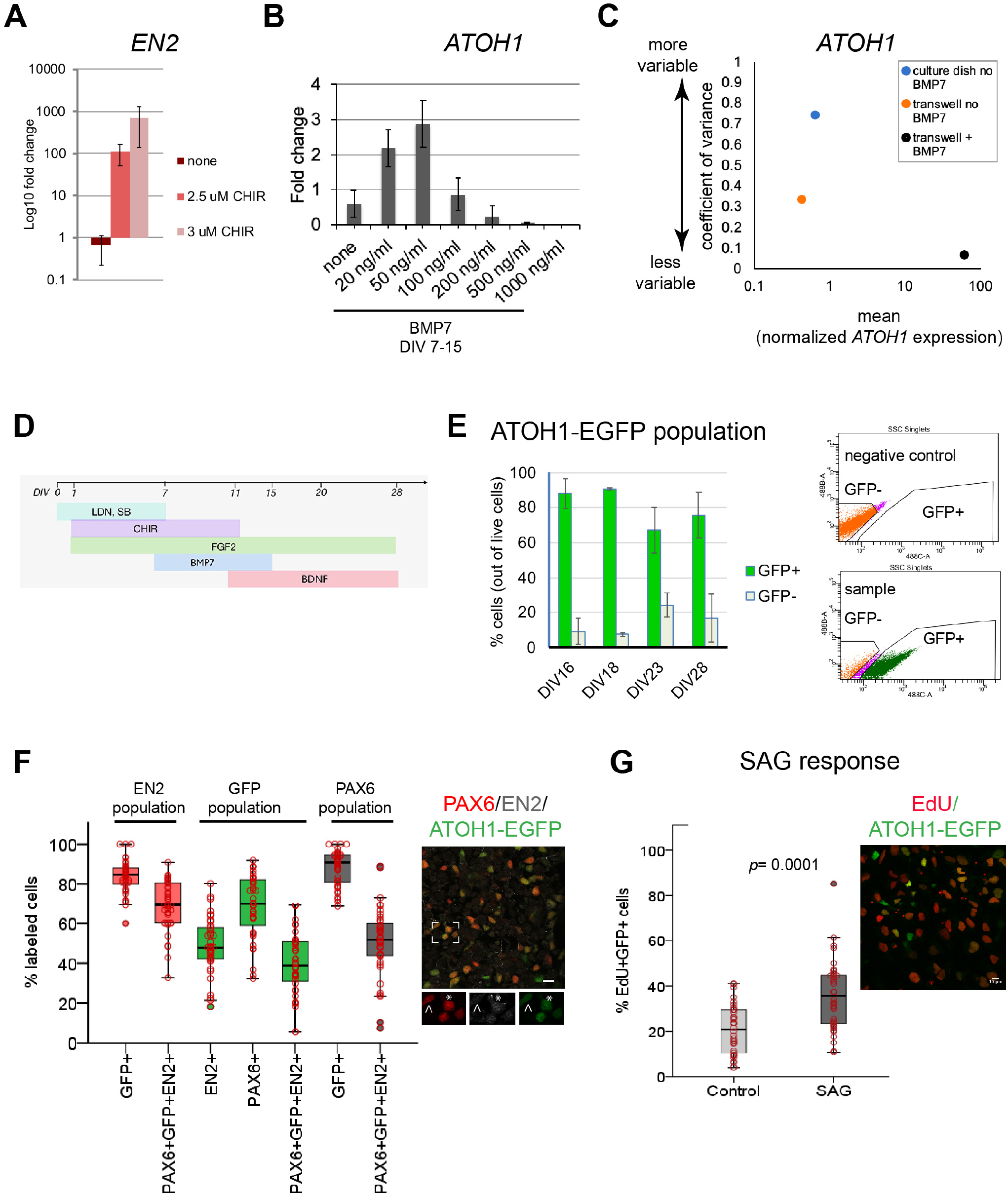
Derivation of the human ATOH1 lineage. **(A)** EN2 expression (fold change of no CHIR) in dual SMAD+FGF2 treated hPSCs in the absence and presence of CHIR99021 by qRT-PCR at DIV11. **(B)** ATOH1 expression (fold change of no BMP7) at DIV16 in response to a BMP7 concentration series added at DIV7-15. **(C)** Dot plot showing the coefficient of variance of mean *ATOH1* expression detected by qRT-PCR at DIV16 in cultures grown on regular tissue culture dishes (-BMP7, blue) versus on transwell membranes (-BMP7, orange; +BMP7, black). **(D)** Schematic of the protocol for derivation of the ATOH1 lineage. (E) Left, the percentage of EGFP+ (green) and EGFP- (grey) cells at DIV16, 18, 23, 28 of differentiation of the *ATOH1-EGFP* line by FAC-sorting. Right, representative FACS charts showing separation of ATOH1-EGFP+/EGFP-cells. **(F)** Left, box plot showing the percentages of EGFP, EN2, PAX6 single and triple positive cells by immunocyto-chemistry, within the EN2, EGPF (ATOH1), and PAX6 populations at DIV28-30. Right, representative merged image of the immunocytochemistry labeling. Boxed area is magnified at the bottom with individual channels displayed. Note asterisk highlighting a triple positive cell, while cell above arrowhead is EN2-;PAX6+;EGFP+. **(G)** Left, boxplot showing the percentage of EdU+;EGFP+ double positive cells per Dapi nuclei +/- SAG treatment after 48 hours (DIV28-30). Right, a representative merged image of the labeling. N= three independent experiments. Bar graphs show mean +/-1 SD Scale bars: 10 μm.

Although this method consistently yielded cerebellar territory in two hPSC lines (Supp Fig. 1C), gene expression levels were variable between experiments. To reduce variability, a common limitation of 2D and 3D stem cell differentiation methods, we tested different culture surfaces (material and area), and the addition of signaling molecules to override stochastic gene expression. Sparse single cell plating of hPSCs on transwell membranes allowed the formation of similarly sized and shaped colonies, resembling the growing neural plate, as patterning proceeded (Supp Fig. 1D). This improved the consistency of the geometric arrangement of the patterned cells as colonies, reduced gene expression variability (Fig. 1C), and increased cell survival, likely due to cells receiving medium in 2D (above and below) compared to 1D in regular culture dishes.

In addition, BMP7 treatment further reduced gene expression variability (Fig. 1C). Previous work has shown that roof plate-derived BMP7 induces *Atoh1* expression in the mouse hindbrain (Alder et al., 1999). Without knowledge of the effective concentration of secreted BMP7 in the developing human, we tested a range of concentrations, and importantly found that low BMP7 concentrations stabilized *ATOH1* expression, while higher levels previously used for mouse ESC-derived GCs (Salero and Hatten, 2007) induced cell death (Fig. 1B). Therefore, the addition of BMP7 appears to override the stochasticity of *ATOH1* expression and reduce variability. Finally, BDNF was added to improve GC survival (Lindholm et al., 1993). Together, these experiments defined optimal conditions for deriving a human cerebellar territory as a first step towards derivation of the ATOH1 lineage and GC differentiation (Fig. 1 D).

To monitor the dynamics of differentiation in culture, we derived a clonal *ATOH1-EGFP* hPSC line, expressing nuclear EGFP under a human *ATOH1* enhancer (Supp Fig. 2A). Fluorescence Activated Cell (FAC)-sorting of ATOH1-EGFP^+^ cells at DIV16 followed by qRT-PCR revealed co-expression of genes associated with the cerebellar territory in mice (Allan Brain Atlas; (Morales and Hatten, 2006) including *ATOH1, PAX6, EN2, ID4, LHX2, LHX9, MEIS2* (Supp Fig. 1E). Time-series analysis of ATOH1-EGFP by imaging and flow cytometry revealed that by DIV16, 80% +/-9 of the cells are ATOH1-EGFP^+^ (Fig. 1E). This is a 5-fold increase in the efficiency of ATOH1^+^ cell derivation compared to a previously reported 3D protocol (80% versus 17%; (Muguruma et al., 2015). By DIV23 the percentage of ATOH1-EGFP^+^ cells decreased, while ATOH1-EGFP^-^ cells increased, indicating the onset of differentiation (Fig. 1E). To investigate when the ATOH1-EGFP^+^ cells co-express known GCP markers, gene expression was analyzed in FAC-sorted ATOH1-EGFP cells at three time-points between DIV16-23. In the mouse, the selected genes are dynamically expressed within the *Atoh1^+^* domain, changing between E11.5, when the *Atoh1^+^* population gives rise to hindbrain and cerebellar nuclei, and E15.5, when the *Atoh1^+^* population forms the EGL (Supp Fig. 1E right, (Machold and Fishell, 2005; Wang et al., 2005); Allan Brain Atlas). Consistent with the reduction in ATOH1-EGFP^+^ cells by DIV23, gene expression shifted between DIV19 and 23, with markers expressed in the mouse *Atoh1* domain at E11.5, prior to EGL establishment (*ID4, LHX2, LHX9*), decreasing in expression, while EGL markers increased (*PAX6, ATOH1*) (Supp Fig. 1E). Thus, at DIV23 the hPSC-ATOH1 lineage initiates GCP production but likely contains a mixture of progenitors. Indeed, at DIV16 a small subset of cells expressed Calretinin by immunohistochemistry (Supp Fig. 2C), a marker of excitatory cerebellar nuclei produced by the Atoh1 lineage prior to GC production. By immunohistochemistry, at DIV 28, the ATOH1^+^ cells co-expressed PAX6 (mean~70%) and EN2 (mean~50%), and ~40% co-expressed all three markers within the GFP^+^ population. The percentage of co-expression of these three markers within the EN2^+^ (marker of mid/hindbrain) population was ~70% (Fig. 1F). Thus, a great majority of the EN2^+^ cells are GCPs. Midbrain progenitors that express EN2 do not express ATOH1/PAX6 (Akazawa et al., 1995) Allan Brain Atlas) and dorsal posterior hindbrain/spinal cord *Atoh1^+^* progenitors, never express EN2 (Davidson et al., 1988).

A defining feature of GCPs is their extensive proliferative capacity in response to sonic hedgehog (SHH) in the postnatal mouse cerebellum (Dahmane and Ruiz i Altaba, 1999; Lewis et al., 2004; Wallace, 1999; Wechsler-Reya and Scott, 1999). EdU uptake, a measure of cell proliferation, was significantly increased in the ATOH1-EGFP^+^ cells (Fig. 1G) after treatment with SAG (an agonist of SHH signaling) compared to control. In conclusion, the hPSC-ATOH1^+^ lineage displayed similar dynamics in gene expression over time as seen in the mouse embryo (albeit extended in period) and produced GCPs, which respond to SAG, by DIV23-28. Compared to previously reported methods, both the speed (35 days versus 16 days) and the yield (from 17% to 80%) of GCP production were increased.

### Differentiation of hPSCs into cerebellar granule cells

In the mouse, early postmitotic GCs switch off *Atoh1* and transiently express TAG1 in the inner EGL. In our cultures, TAG1 was expressed already at DIV18, after which its expression increased (FIG. 2A). Magnetic Activated Cell Sorting (MACS) using an antibody against TAG1, which is expressed on the cell surface, yielded 13% +/-7 (n=7) TAG1^+^ cells at DIV28 (Fig. 2 B,C). TAG1^+^ cells were then co-cultured with either mixed cerebellar mouse neurons or glia to provide a permissive differentiation environment. By DIV3 in co-culture, the majority of the TAG1^+^ cells were PAX6^+^ (Fig. 2D, top panel). By contrast, the TAG1^-^ cells had larger nuclei and were PAX6^-^ (Fig. 2D, bottom panel). By DIV20 in co-culture (DIV48 total days), a majority of the human cells displayed characteristic GC morphology, namely a small (<10 μm) round nucleus and a limited number of extended processes (3-4) including bifurcated axons. Mouse GCs isolated at P0 and grown concurrently in the same dish as the human cells displayed similar features (Fig. 2E). The human cells also expressed the GC markers NEUROD1 and NeuN (Fig. 2E and Supp Fig. 2C, *bottom panel*). Interestingly, pre-synaptic specializations were apparent in neurons grown in co-culture with mouse glia only, indicating that cross-species neuron-glia interactions are sufficient to trigger synapse initiation (Fig. 2F).

**Figure 2.**
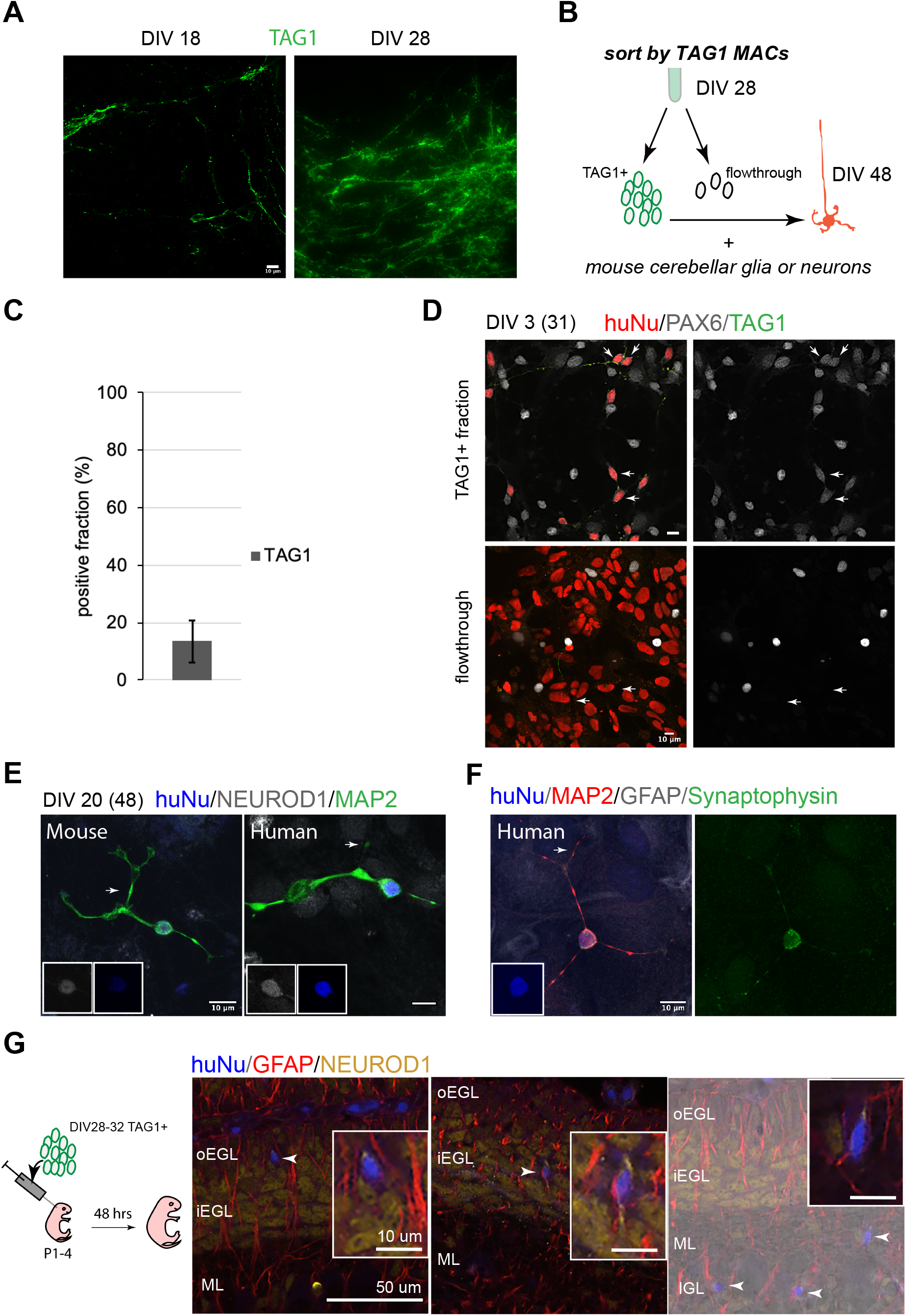
Human GC differentiation in co-culture and integration into the mouse cerebellum upon transplantation. **(A)** TAG1 expression at DIV18 (left) and DIV28 (right). **(B)** Schematic of the sorting strategy of TAG1+ cells by MACS at DIV28 and co-culture with mouse cerebellar neurons or glia until DIV48. **(C)** Bar chart (mean+/- 1 SD) of TAG1+ cells/ total cells at DIV28, N=7 experiments. **(D)** *Top*, TAG1+ cells (green) in co-culture with mouse cerebellar neurons and glia for three days express PAX6 (red+white, arrows). *Bottom*, TAG1-cells (flowthrough) in co-culture with mouse neurons and glia have larger nuclei and are PAX6-(arrows). **(E)** TAG1+ cells in co-culture with mouse neurons and glia for 20 days (DIV48 total), display small round nuclei (inset, blue), bifurcated neuronal extensions, and are NEUROD1 +;Map2+ (see F). A mouse GC cultured in the same dish for the same period of time, shown for comparison. **(F)** TAG1 + cell in co-culture with glia only for 20 days (DIV48 total) expresses Synaptophysin. **(G)** Three representative coronal sections of mouse cerebella transplanted with day 28-32 TAG1+ human cells, 48 hrs post transplantation. Far right image is overlaid on a DIC image. Boxed images are higher magnifications of migrating cells. HuNu, human nuclear antigen; IGL, internal granule cell layer; oEGL, outer EGL; iEGL, inner EGL; ML, molecular layer. Scale bars as indicated on images.

To assess whether hPSC-GCs would integrate into the mouse cerebellar cortex, especially whether they would undergo the classic glial-guided migration of GCs, we implanted human cells into the juvenile mouse cerebellum at the time when mouse GCs are undergoing migration. Importantly, TAG1 sorted cells integrated into the neonatal mouse cerebellar cortex and migrated along glial fibers with stereotypical morphology of migrating neurons passaging through the EGL to settle in the IGL (Fig. 2G).

### hPSC-ATOH1^+^ cells match the molecular profile of the human cerebellum in the second trimester

To transcriptionally profile the hPSC-ATOH1^+^ progenitors, we adapted the TRAP methodology, first developed in transgenic mice for cell-type specific translational profiling (Doyle et al., 2008; Heiman et al., 2008), for hPSCs. An EGFP-tagged L10a ribosomal subunit was driven by the human *ATOH1* enhancer (Supp Fig. 2B), enabling GFP-mediated immunoprecipitation (IP) of ribosomally attached mRNAs in the ATOH1 lineage specifically. Importantly, this method bypasses the need for cell dissociation and provides information about transcripts, including the 3’UTR, that are present in both the cell soma and processes. TRAP followed by RNA sequencing (TRAP-seq) was performed at DIV28, when the ATOH1 lineage co-expressed GCP markers and displayed increased proliferation in response to SAG (Fig. 1). The *ATOH1* transcript was indeed enriched in IPs compared to the input (Supp Fig. 2B, 3A). Several other GCP markers were similarly enriched, while markers of differentiated GCs were depleted (FIG. 3A). Heatmaps of the major signaling pathways in development (WNT, BMP, SHH, FGF, HIPPO) highlighted enrichment of the WNT pathway in particular (Supp Fig. 3B). Interestingly, gene ontology analysis revealed axon guidance, WNT pathway, neuronal migration, and cell division among the top significantly enriched developmental processes (Supp Table1).

**Figure 3.**
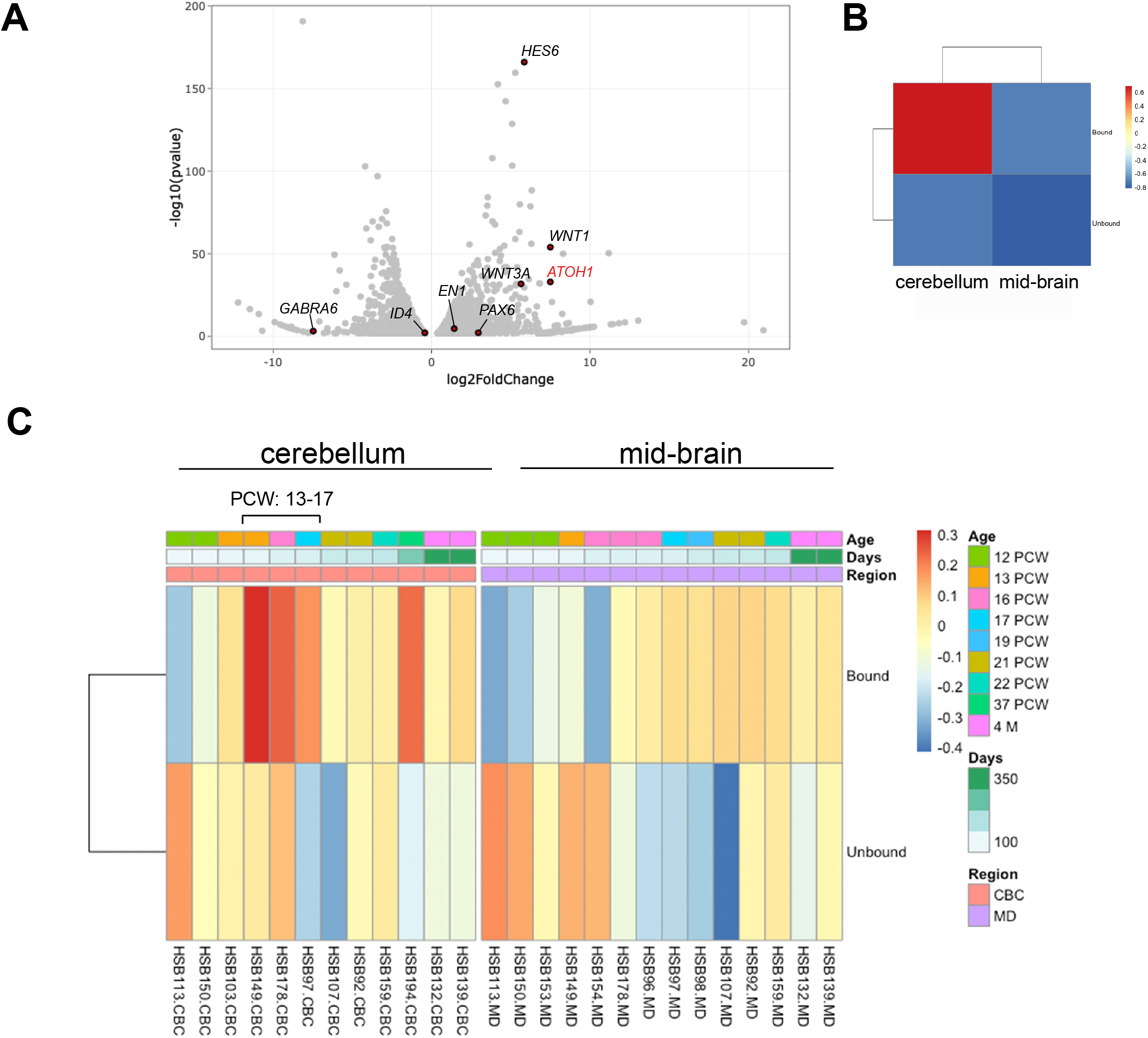
The hPSC-derived ATOH1 lineage resembles the human cerebellum in the second trimester by translational profiling. **(A)** Volcano plot of Log2 fold change global gene expression in ATOH1-TRAP IPs versus input. Key GC genes are highlighted in red. The fully differentiated GC marker GABRA6 is depleted while progenitor genes are enriched. **(B)** Heatmap showing GSEA analysis of Log2 fold enriched genes in DIV28 ATOH1-TRAP versus the PsychEncode dataset for the developing human cerebellum and midbrain from 12 PCW until 4 months of age (combined). **(C)** Heat map of data in B but divided by timeline with columns representing our data (bound (IP) and unbound (input)) compared to individuals from the PsychEncode project (identifiers depicted at the bottom). CBC, cerebellum; MB, midbrain; PCW, post coitus weeks.

To investigate how the hPSC-ATOH1^+^ cells compare to the molecular signature of the developing human cerebellum, we compared our dataset to RNA-seq data from the PsychEncode study (Li et al., 2018), which samples different brain regions during a developmental timeline covering 8 post coitus week (PCW) until after birth in the human. Gene Set Enrichment Analysis (GSEA) revealed that the DIV28 ATOH1 lineage most closely matched the profile of the 13-17 PCW human cerebellum (Fig. 3 B, C). Together, these analyses provide the first translational data set for the hPSC-derived ATOH1 lineage. While the data closely match the developing human cerebellum, development appears accelerated in culture.

### Translational profiling reveals a transcriptional heterochronic shift in the human versus mouse cerebellum

In a surprising finding, TRAP-seq analysis of the *ATOH1* lineage revealed the expression of several genes that are classically associated with postmitotic GCs in mice including *RBFOX3* (encodes for the NeuN antigen) and *NEUROD1* (Fig. 4A). The co-expression of *NEUROD1* and *ATOH1* was confirmed by RT-PCR in IPs (Fig. 4A) and in FAC-sorted EGFP^+^ cells derived from the *ATOH1-EGFP* line (data not shown). To investigate if the expression of these factors extends into the proliferative zone of the human EGL *in vivo* or if this is an *in vitro* phenomenon, we performed immunohistochemistry in the human cerebellum at 17 PCW. In contrast to the mouse, many outer EGL cells, where the *ATOH1^+^* progenitors reside (Haldipur et al., 2019), express both NEUROD1 (Fig. 4B, D, Supp. Fig. 4) and NeuN (Fig. 4C, D). Moreover, Ki67, a marker of proliferating cells, was not as prevalent at this stage in the human EGL as it is in a comparable stage of cerebellar development in the mouse (based on foliation depth and pattern, P0, (Biran et al., 2012; Haldipur et al., 2019) (Fig. 4B-high mag and C). In the human EGL, Ki67 expression was sparser, Ki67 and NEUROD1 co-expression was clearly evident, and there was no clear separation of cells into two layers as seen in the mouse. Punctate NeuN expression was detected in the human oEGL but not in the mouse (Fig. 4D). Together, these data reveal extensive co-expression of proteins that have classically until now been ascribed to either progenitors or differentiating neurons based on mouse studies.

**Figure 4.**
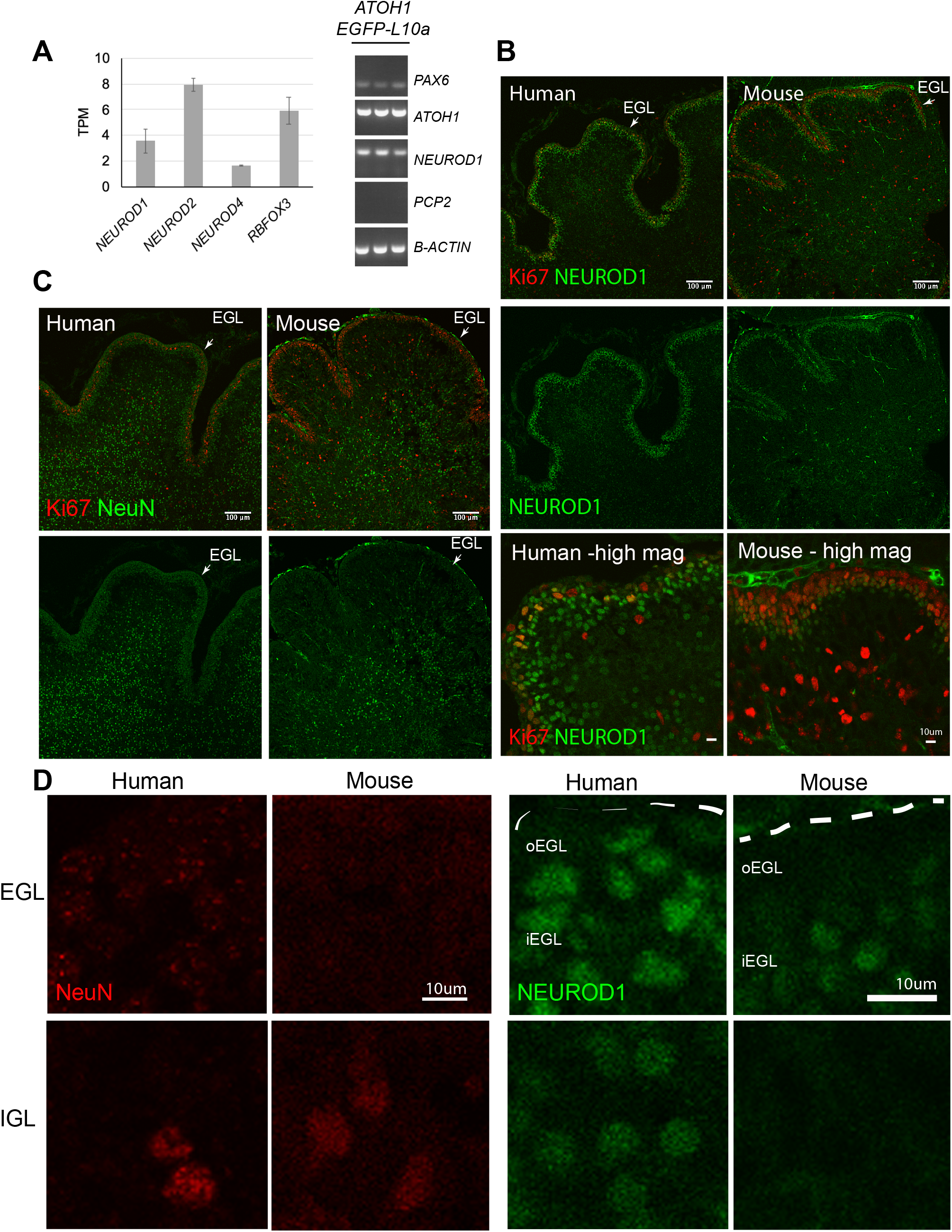
Heterochronic shift in the expression of transcriptional regulators in the human EGL compared to mouse. **(A)** *Left*, bar chart showing the mean normalized expression of transcriptional regulators in ATOH1-TRAP IPs at DIV28 by RNA-seq. *Right*, The expression of *PAX6* (GCP marker), and co-expression of *ATOH1* and *NEUROD1,* but not *PCP2* (Purkinje cells marker) in ATOH1-TRAP IPs by RT-PCR. **(B)** Sagittal sections though the vermis showing NEUROD1 and Ki67 expression by immunohistochemistry in the human cerebellum at 17 PCW and mouse at P0. Note the similarities in foliation depth and pattern. *Bottom*, higher magnification (scale bars: 10 μm) of a lobule in human and mouse. **(C)** Mid-sagittal sections showing NeuN (RBFOX3) and Ki67 in the human (17 PCW) and mouse (P0). **(D)** Higher magnification of NeuN and NEUROD1 labeling in the human versus mouse EGL and IGL (scale bars: 10 μm). *Left* panel, note the punctate NeuN labeling in the human but not the mouse EGL. *Right* panel, NEUROD1, dashed lines demarcate the pial surface. EGL, external granule cell layer; IGL, internal granule cell layer; TPM, transcripts per million reads. Scale bars: 100 μm unless stated otherwise.

## Discussion

We report a method for the scalable derivation of the human ATOH1 neuronal lineage from hPSCs that yields cerebellar progenitors by day 16 and GCs within 48 days in chemically defined medium. In contrast to previous methods for derivation of cerebellar neurons with a focus on Purkinje cells (Buchholz et al., 2020; Erceg et al., 2010; Muguruma et al., 2015; Nayler et al., 2017; Silva et al., 2020; Wang et al., 2015; Watson et al., 2018), we provide a timeline of developmental progression of the human ATOH1 lineage by gene expression and an unbiased comparison of the transcriptional profile to the developing human brain. While our focus was derivation of GCs, the expression of markers of the cerebellar nuclei (LHX9, LHX2) and the presence of Calretinin^+^ cells indicate the production of additional ATOH1-derivatives. Future identification of cell surface markers for each class of glutamatergic neuron known to derive from the ATOH1 lineage will allow isolation and further characterization of the various populations yielded by our method. MAC-sorted early postmitotic GCs (TAG1^+^) integrated into the mouse cerebellum and migrated through the EGL into the IGL in close apposition to glial fibers, providing an *in vivo* system to study human GC migration and migration defects. This is the first successful demonstration of migration followed by integration of hPSC-GCs into a stratified cortex, suggesting that molecular pathways required for glial-guided migration have developed in the human cells.

Adaptation of TRAP-seq for use in hPSCs allowed lineage-specific molecular profiling of the ATOH1 lineage, providing a superior depth of read compared to scRNA-seq methods. Comparison of our data to RNA-seq data from the developing human brain, matched the DIV28 hPSC-ATOH1 lineage to the human cerebellum at 13-17 PCW. Analysis of key developmental pathways revealed that the WNT pathway is particularly enriched. Indeed, mouse genetic studies show that WNT signaling is critical for both early cerebellar development and later circuit establishment (Lucas and Salinas, 1997; McMahon and Bradley, 1990).

Interestingly, using TRAP-seq methodology we identified a heterochronic shift in the expression of transcriptional regulators in progenitors in the human EGL, that in the mouse are expressed in cells that are further along the differentiation path. Genetic differences in developmental timing can cause birth defects or give rise to a novel morphology that could confer an evolutionary advantage. The temporal shift in NEUROD1 and RBFOX3 expression in the human oEGL suggests an expansion of an “intermediate” cellular state that may serve to allow the progenitor pool to persist much longer, to enable the protracted period of cerebellar development in humans compared to mouse. Both NEUROD1 and RBFOX3 are expressed in early postmitotic as well as fully differentiated neurons, rarely in neuroblasts, across species and brain regions (D’Amico et al., 2013; Lee et al., 1995; Miyata et al., 1999; Zhang et al., 2016). Overexpression of either gene in progenitors induces neuronal differentiation (Boutin et al., 2010; Butts et al., 2014; Lee et al., 1995; Pataskar et al., 2016; Zhang et al., 2016). Interestingly, in the *Xenopus* embryo, the co-expression of *Atoh1* and *NeuroD1* in the EGL has been reported and it was hypothesized that both the timing of expression and the gene regulatory function of NeuroD1 have adapted to support development of a non-proliferative EGL (Butts et al., 2014). In the human, we propose that the extensive co-expression of these factors may enable a quiescent state, to preserve the progenitor pool. Indeed, Ki67 labeling showed fewer positive cells in the human EGL at 17 PCW than the comparable P0 mouse EGL, hinting at differences in proliferative regulation. Our findings, together with findings in the *Xenopus* and the differential proliferative capacity of the EGL in several vertebrate species (Iulianella et al., 2019), support the existence of evolutionary adaptations in EGL development across species. In future work, analyses of cell cycle length and comparison of molecular features in human versus mouse GCPs should provide further insights into the molecular basis of human cerebellar expansion and human-specific molecular mechanisms that will be critical for understanding human medulloblastoma pathogenesis as well as cerebellar-mediated human cognitive evolution.

---

**Supplementary Figure 1.**
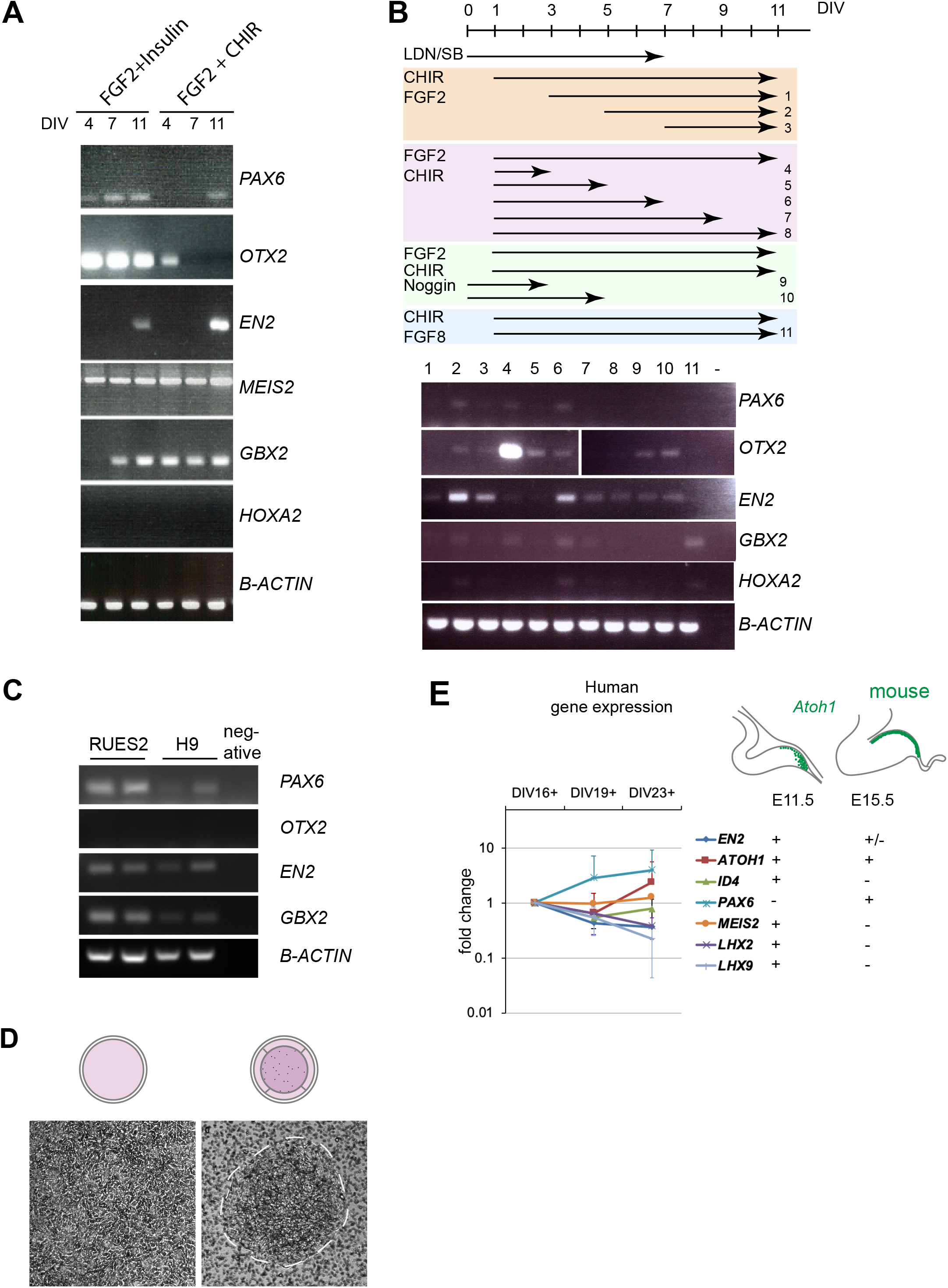
Derivation and characterization of the human ATOH1 lineage. **(A)** Gene expression by RT-PCR at DIV4, 7, and 11 comparing dual SMAD inhibition plus FGF2/Insulin versus FGF2/CHIR99021 treatment. **(B)** *Top*, schematic showing 11 different time intervals of FGF2, CHIR99021, and Noggin treatment in the presence of dual SMAD inhibition at DIV0-7 versus FGF8/CHIR99021. *Bottom*, RT-PCR at DIV11 showing the resulting expression of various markers in the 11 different conditions. **(C)** RT-PCR of granule cell progenitor markers at DIV16 in RUES2 and H9 hESC lines upon dual SMAD inhibition (DIV0-7) plus FGF2/CHIR99021 (DIV1-11) treatment. **(D)** Comparison of differentiating cells on regular culture plates (left) versus transwell plates (right). Dashed lines demarcate the edge of a colony. **(E)** Right, schematic showing how gene expression in the mouse Atoh1 lineage changes from embryonic day (E)11.5, prior to EGL establishment, and at E15.5 in the EGL. + indicates presence of expression, - indicates absence of expression. *Left*, qRT-PCR detection of gene expression of listed genes in ATOH1-EGFP FAC-sorted cells at DIV16, 19, 23 of culture. All genes were detected at DIV16 and fold change is relative to levels at DIV16. N=3 independent experiments. DIV, days in vitro; LDN, LDN193189; SB, SB431542.

**Supplementary Figure 2.**
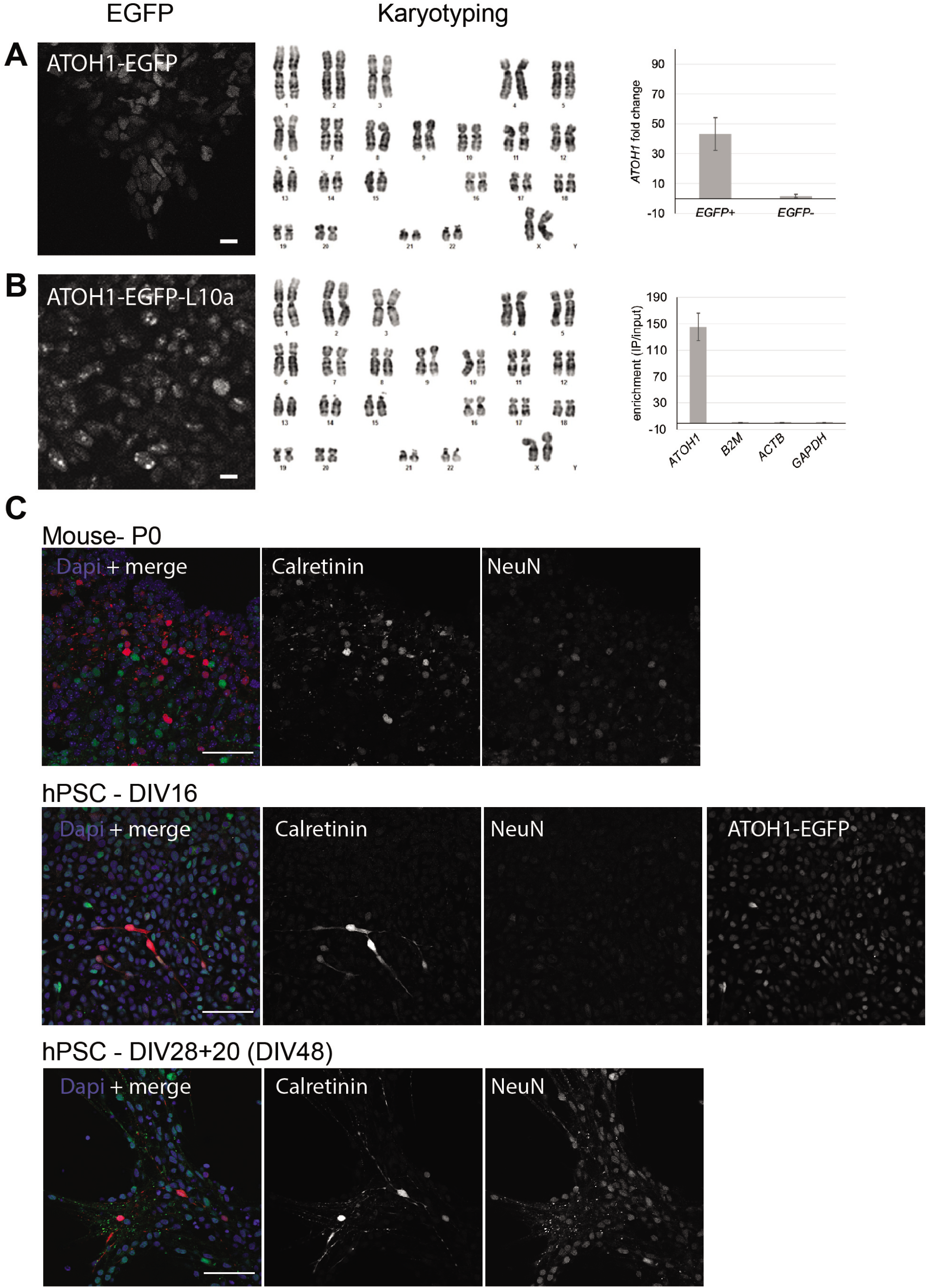
Characterization and differentiation of ATOH1-EGFP and ATOH1-EGFP-L10a transgenic lines. **(A)** Left, Representative image of EGFP expression upon differentiation of the *ATOH1-EGFP* lines at DIV16. Middle, normal karyotype detected. Right, bar chart showing the mean +/- SD of normalized *ATOH1* expression levels in FAC-sorted EGFP+ cells as fold change of expression in EGFP-cells at DIV28 by qRT-PCR. **(B)** Left, Representative image of EGFP-L10a expression upon differentiation of the *ATOH1-EGFP-L 10a* lines at DIV16. Note the marked difference in the localization of GFP compared to A. Middle, normal karyotype detected. Right, bar chart showing the mean +/- SD of the level of enrichment of *ATOH1* compared to three house-keeping genes in *ATOH1-EGFP-LWa* TRAP IPs versus input at DIV28 by RNAseq. **(C)** Top panel, Calretinin and NeuN expression in the cerebellar nuclei at P0 in mice. Middle, Calretinin, NeuN, and ATOH1-EGFP expression in hPSC cultures detected at DIV16. Bottom, Calretinin and NeuN detected at DIV48. Note that NeuN expression increases with time in culture and like in mice, most Calretinin neurons are NeuN negative.

**Supplementary Figure 3.**
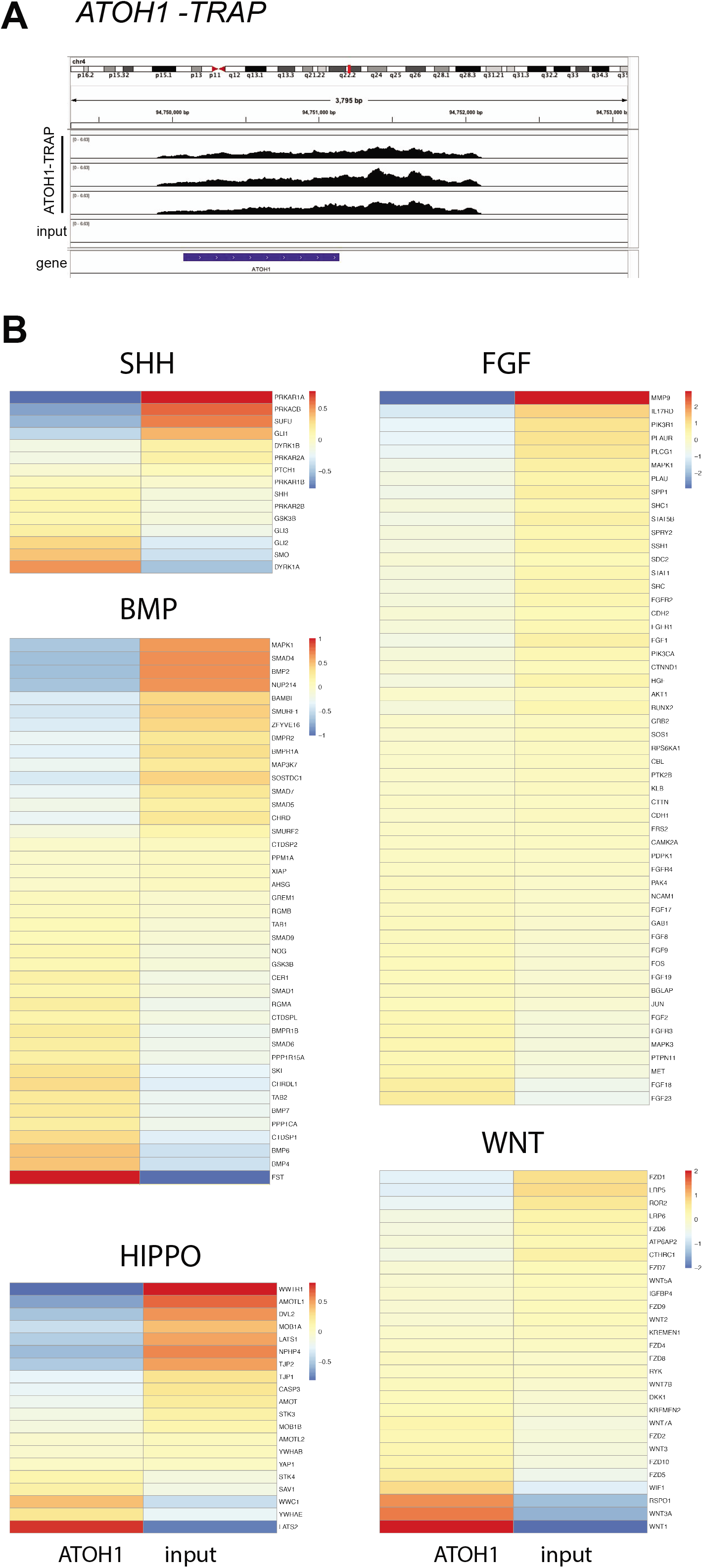
Heatmaps of key developmental signaling pathways in the hPSC-ATOH1 lineage. **(A)** RNA sequencing of immunoprecipitated mRNAs from an *ATOH1-EGFP-L10A* hPSC TRAP line differentiated until DIV28 shows enrichment of *ATOH1* reads in the ATOH1-TRAP IPs compared to the input, depicted by Integrative Genomics Viewer. **(B)** Heatmaps showing the enrichment of genes in the *ATOH1-EGFP-L10a* TRAP versus input. Genes have been organized according to the developmental pathways they are associated with including the SHH, BMP, HIPPO, FGF, and WNT pathways.

**Supplementary Figure 4.**
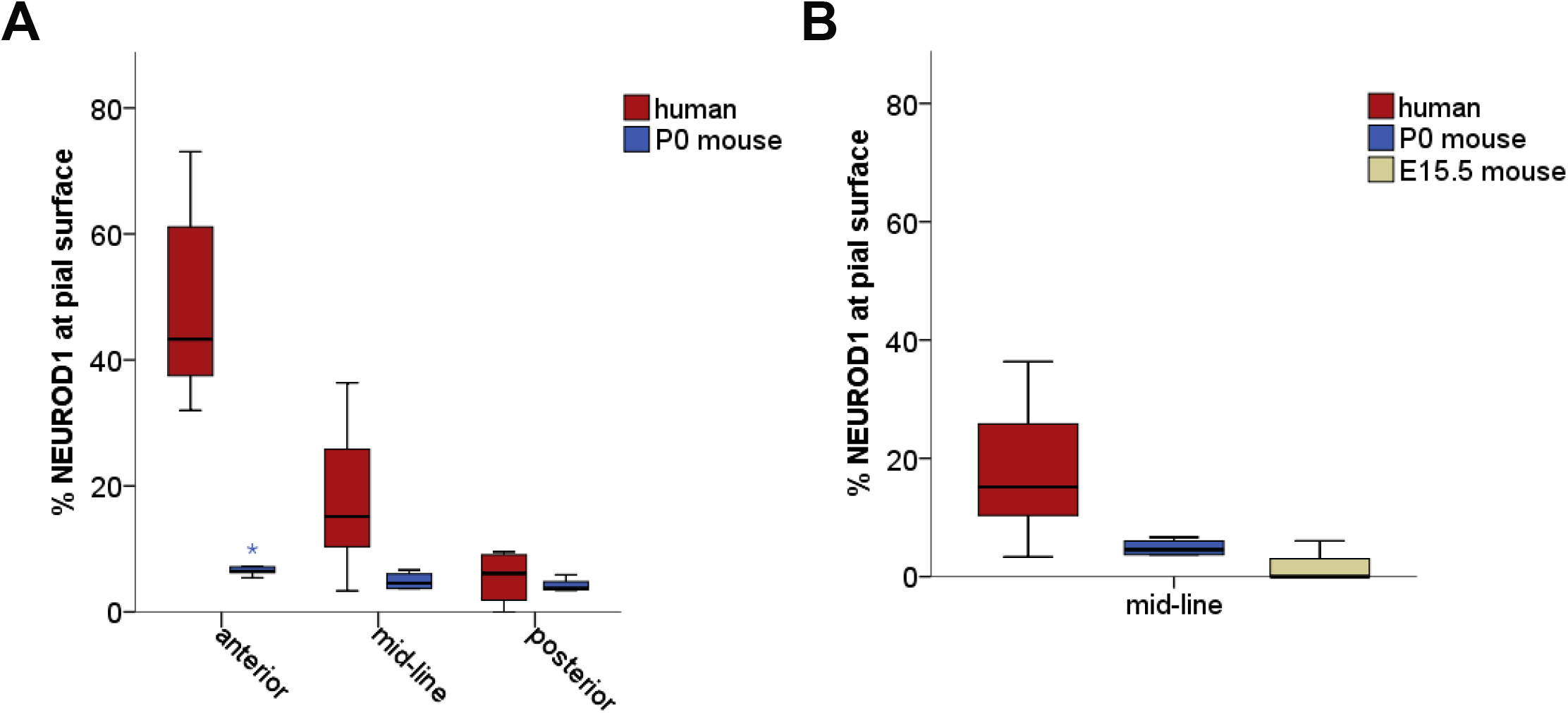
Quantification of NEUROD1 expression at the pial surface of the developing cerebellum in the human compared to mouse. **(A)** Box plot showing the percentages of NEUROD1+ nuclei per total nuclei (Dapi) along the length of the pial surface in anterior, mid-line, and posterior regions of the developing cerebellum comparing mouse (P0) to human (17 PCW). Note that the percentage of NEUROD1+ cells at the pia is low across the anterior-posterior axis of the mouse cerebellum, in contrast to the human where the anterior-posterior axis differs (Human n. cells at the pia counted = 613, Human NEUROD1+ fraction anterior: 48.4 +/- 6.5 SE, mid-line: 21.1 +/- 4.3 SE, posterior: 5.4 +/- 2.2 SE. P0 Mouse n. cells at the pia counted = 369, P0 Mouse NEUROD1+ fraction anterior: 7.1 +/- 0.8 SE, mid-line: 4.8 +/- 0.7 SE, posterior: 4.3 +/- 0.8 SE). **(B)** For comparative purposes an additional earlier (E15.5) stage of development was analyzed in the mouse. Box plot showing data from mid-line regions of the cerebellum in 17 PCW human, P0 mouse, and E15.5 mouse cerebellum. Few NEUROD1+ cells were detected at the pial surface in the mouse at multiple stages of development in contrast to the human (E15.5 Mouse n. cells at the pia counted: 188, NEUROD1+ fraction mid-line: 2 +/- 2 SE, posterior: 1.8 +/- 1.8 SE).

## Methods

### Human tissue collection, fixation, and embedding

Human prenatal brain tissue (2 cerebella; 17 PCW) were acquired from the Human Developmental Biology Resource (http://www.hdbr.org/) following institutional policies. Tissue were fixed in 4% PFA for 7-10 days and washed multiple times in PBS. Samples were then cryoprotected in increasing concentrations of 5%, 15%, and 30% sucrose (in PBS) at 4°C. Samples were embedded in Tissue-Tek O.C.T. compound (VWR, 25608-930) and stored at −80°C until use.

### Mice

All procedures with mice were performed according to guidelines approved by the Rockefeller University Institutional Animal Care and Use Committee. C57Bl/6J mice (Jackson Laboratory) were maintained on a 12 hr light/dark cycle and a regular diet. Timed matings generated a mixture of male and female pups that were used for all described studies.

### hPSC culture

Human embryonic/pluripotent stem cells were used under approved institutional ESCRO committee protocols (The Rockefeller University). The RUES2 and H9 (WA09) hESC lines (a kind gift from Dr. Ali Brivanlou, The Rockefeller University) were maintained in growth media (HUESM medium conditioned with mouse embryonic fibroblasts and supplemented with 20ng/ml bFGF (Invitrogen), (Deglincerti et al., 2016). The growth medium was exchanged daily. For transgenic lines, Puromycin (Gibco) was added to the medium (1μg/ml) during maintenance culture. Cells were grown as colonies on tissue culture dishes coated with hESC qualified Matrigel solution (Corning) in a 37°C humidified incubator with 5% CO^2^.

### hPSC differentiation

hPSCs maintained as colonies were dissociated from plates with Trypsin-EDTA (0.25%, Gibco) for 4 minutes at 37°C in a humidified incubator. Cells were washed once with growth media (see previous section) and then re-suspended in growth media with 10 μM ROCK-inhibitor Y-27632 (Abcam). Single cells were plated at 900 cells/ml on Transwell 6-well plates with permeable 24 mm polyester membrane inserts (Corning, 3450) or regular tissue culture-treated plates. On Transwell dishes, cells were plated on top of the membrane with 1ml growth medium plus ROCK-inhibitor added below and above the membrane (2 ml total). The next day, 1 ml growth medium plus ROCK-inhibitor was added to the top part of the Transwell (3 ml total). On day two, the medium was switched to differentiation medium: DMEM/F12 with sodium bicarbonate (Invitrogen, 11320-033), 0.05% BSA, 0.1 mM β-mercaptoethanol, 2 mM Glutamate, 10 μM NEAA, 1x N2 supplement, 1x B27 without retinoic acid (all from Gibco) and Primocin (0.1 mg/ml, InvivoGen). This marked day 0 of differentiation. For differentiation experiments with transgenic lines, Puromycin (Gibco) was added to the medium (0.5 μg/ml) until DIV28. The following small molecules and growth factors were added to the differentiation medium on days indicated in Fig. 1D as follows: 10 μM SB431542 (SB, Tocris) and 100 nM LDN-193189 (Stemgent) at days 0-7, 2.5 μM CHIR99021 (Stemgent) at days 1-11, 20 ng/ml bFGF (Invitrogen) at days 1-28, 25 μg/ml BDNF at days 11-28, 50 ng/ml recombinant mouse BMP7 (R&D Systems) at days 7-15. Medium was exchanged for fresh differentiation medium plus appropriate factors every other day until day 28. The following conditions were also tested (not all data were included in the manuscript) to arrive to the optimized protocol for the derivation of the ATOH1 lineage: CHIR99021 (Stemgent) at days 1-11 (range tested: 1-3 μM), 7 μg/ml human recombinant Insulin (Tocris), 100 ng/ml recombinant human/mouse FGF8b (Tocris) between days 1-11 (range tested: 50-500 ng/ml), recombinant mouse BMP7 (R&D Systems) at days 7-15 (range tested: 20-1000 ng/ml), 20 ng/ml BMP6 (R&D Systems), BMP4 (range tested: 4-160 ng/ml, R&D Systems), 100 ng/ml GDF7 (R&D Systems). Depending on culture conditions used, such as culture surface, the concentration of CHIR99021 may need to be adjusted to achieve optimal anterior hindbrain patterning.

### Generation of transgenic hESC lines

For derivation of the *ATOH1-EGFP* line, a human *ATOH1* enhancer sequence (GenBank Accession number AF218259.1, (Helms et al., 2000) was amplified from human genomic DNA using two infusion primers (forward primer: 5’-TTCAAAATTTTATCGATAAGGTTCTTCTATGGAGTTTGCA-3’, reverse primer: 5’-AATAGGGCCCTCTAGAGAATTCCTGAACAACCCCAC-3’). The amplicon was cloned into a modified version of the self-inactivating lentiviral vector pPS-EF1α-GFP-RFP (System Biosciences, LV603PA-1) where GFP and RFP had been removed and replaced by an hPGK-Puromycin cassette (pSIN-EF1a promoter-BGH polyA-hPGK-Puromycin). The vector was digested with *Clal* and *Xbal* enzymes to remove the *EF1a* promoter, and the *ATOH1* enhancer sequence was cloned using in-fusion HD cloning (Clontech). Subsequently, a sequence containing the human beta globin minimal promoter (*hβ-globin*) followed by a nuclear localization signal and *EGFP* (nls-EGFP) was amplified from the J2XnGFP plasmid DNA (a gift from Dr. Jane Johnson, UT Southwestern) and cloned downstream of the human *ATOH1* enhancer by in-fusion HD cloning with the following primers (nls-EFGFP forward primer: 5’-GGTTGTTCAGGAATTCGATGGGCTGGGCATAAAAGT-3’, nls-EFGFP reverse primer: 5’-GCCCTCTAGAGAATTCAACTAGAGGCACAGTCGAGGC-3’) to obtain the following lentiviral construct: pSIN-hATOH1 enhancer-hβ-globin-nlsEGFP-bGH polyA-hPGK-Puromycin. For derivation of the *ATOH1-EGFP-L10a* line, the *nlsEGFP* was replaced with *EGFP-L10a*. Briefly, the pSIN-hATOH1 enhancer-hβ-globin-nlsEGFP-bGH polyA-hPGK-Puromycin construct was digested with *SbfI/BsrGI* enzymes to remove the *nlsEGFP*. An *EGFP-L10a* fusion fragment was amplified from a template plasmid (mPCP2-A box-s296, a gift from Dr. Nathaniel Heintz, The Rockefeller University) with the following in-fusion PCR primers (EGFP-L10a-forward primer: 5’-CATTTGCTTCTAGCCTGCAGGTCGCCACCATGGTGAG-3’ and EGFP-L0a reverse primer: 5’-CCGCTTTACTTGTACATTATCTAGATCCGGTGGATCC-3’) and cloned by in-fusion HD cloning to obtain pSIN-hATOH1 enhancer-EGFP-L10a-bGH polyA-hPGK-Puromycin. Lentiviral mediated delivery was used according to established protocols to introduce the vectors into the RUES2 hESC line to generate transgenic lines. Two clonal lines with normal karyotype (Molecular Cytogenetics Core facility, MSKCC) and pluripotency characteristics were selected and expanded per transgenic line. Upon differentiation, EGFP expression was examined by microscopy. In the *ATOH1-EGFP* line, EGFP was broadly expressed in the nuclei of a subset of cells, while in the *ATOH1-EGFP-L10a* line, EGFP puncta were localized to the nucleoli, the site of ribosomal biogenesis (Supp. Fig. 2A, B). A majority of the labeled cells co-expressed Ki67 (proliferation marker, data not shown), and the *ATOH1* transcript was enriched in FAC-sorted EGFP^+^ cells from the *ATOH1-EGFP* line compared to EGFP^-^ cells, and upon IP in the *ATOH1-EGFP-L10a* line compared to input. By contrast, housekeeping genes were not enriched (Supp. Fig. 2A, B).

### Fluorescence Activated Cell Sorting (FACS)

Cells at days 16, 19, 23, and 28 of differentiation were FAC-sorted on a BD FACSAriaII with BD FACSDiva 8.0.1 software (BD Biosciences), using a 100 μm nozzle and a 488 nm laser according to standard procedure. Briefly, differentiation cultures were washed once in Ca^2+^/Mg^2+^ free PBS and the cells were dissociated by incubation in Accutase (Millipore, SCR005) for 5 min at 37°C. Dissociated cells were resuspended in MACS buffer (see MACS sections) and put through a cell strainer (BD Falcon, 352235). Gating was performed on EGFP positive and negative control cells and Propidium Iodide (Sigma), at an appropriate dilution, was used for dead cell exclusion. EGFP^+^ cells were collected in MACS buffer for further downstream analyses.

### Magnetic Activated Cell Sorting (MACS) of TAG1^+^ cells

On days 28-32 of differentiation, TAG1^+^ cells were isolated by MACS (Miltenyi Biotec) according to the manufacturer’s protocol. Briefly, cells were washed 1x in MACS buffer (0.5% BSA, 0.9% glucose in Ca^2+^/Mg^2+^ free PBS) and then gently scraped off from Transwell membranes in MACS buffer, using a cell scraper (USA Scientific, CC7600-0220). Cells were collected by centrifugation (300 g, 10 min). 1 ml Trypsin (1 g/ml)-DNase (100 mg/ml) solution (Worthington Biochemical, 3703 and 2139) was added to the pellet for 1.5 min at 37°C without disturbing the pellet. The Trypsin-DNase was then exchanged for 1 ml DNase (100 mg/ml) and the cell pellet was triturated using a fine-bore pulled glass pipette until a uniform cell suspension was obtained. The suspension was put through a 40 μm cell strainer (BD Biosciences, 352340) to remove remaining clumps. Serum containing “cerebellum” media (see subsequent section, plus 10% horse serum, Invitrogen 26050-088) was added to inactivate the Trypsin-DNase and the single cell suspension was washed and spun (300 g, 10 min) in 50 ml of CMF-PBS (PBS ThermoFisher, 14190-250; 0.2% w/v glucose Millipore, G8769; 0.004% v/v NAHCO_3_ Millipore, S8761; 0.00025% Phenol Red Millipore, P0290). The cells were resuspended in 1 ml serum containing “cerebellum” medium and spun in a tabletop centrifuge (300 g, 5 min) in an Eppendorf tube. This step helped reduce dead cells and debris, which accumulated in the supernatant. The cell pellet was then resuspended in fresh “cerebellum” medium plus serum and incubated in a bacterial dish in a 35°C, 5% CO_2_ incubator for 1 hr. This step allowed surface antigens to reappear after the enzymatic dissociation of cells. The number of live cells was counted and the cells were incubated in TAG1 antibody in cerebellum medium (1:2) for 20 min at room temperature (RT), followed by 1x wash in MACS buffer, and then incubated in anti-mouse IgM microbeads (Miltenyi Biotec, 130-047-302) for 15 min at RT and TAG1^+^ cells were sorted through MS columns (Miltenyi Biotec, 130-042-201) following the manufacturer’s description.

### Purification of cerebellar neurons and glia and co-culture with human cells

Mixed cerebellar cultures or glial fractions were prepared from P0-1 pups and cultured in serum-free “cerebellum” media (BME, Gibco; 2mM L-Glutamate, Gibco; 1% v/v BSA, Sigma; ITS liquid media supplement, Sigma; 0.9% v/v glucose, Sigma; 0.1mg/ml Primocin, InvivoGen) as previously described (Baptista et al., 1994; Hatten, 1985). Mixed cerebellar cultures (no separation step) were plated at 2.8 x 10^6^ cells/ml on poly-D-lysine (Millipore) coated glass coverslips (Fisher, 12545-81) placed in 24-well plates. Glia-only fractions (separated by Percoll gradient) were plated at 0.8 x 10^6^ cells/ml. TAG1^+^ human cells isolated at DIV28 by MACS were then plated on top of mixed cerebellar cultures the next day (at 0.3-0.5 x10^6^ cells/ml). Glial cultures were allowed to form a monolayer (5-7 days) upon which DIV28 TAG1^+^ cells were then plated. As previously reported for their mouse counterparts (Hatten, 1985), the ratio of glia to human neurons in co-culture was crucial. At 1:4 (glia:neuron), the neurons induced detachment of glia from the culture dish and co-aggregated into attached spheres while at a 1:2 (glia:neuron) ratio, the glia remained as a bed upon which neurons attached and extended processes. Half of the medium (serum-free cerebellum media) was replaced with fresh medium every 4 days.

### Gene expression analysis by RT-PCR and qRT-PCR

mRNAs were extracted using the RNeasy Plus mini kit (Qiagen) with on column genomic DNA elimination, and cDNAs transcribed with the Transcription First Strand cDNA Synthesis Kit (Roche) according to the manufacturer’s description. Reverse transcription and PCR were carried out according to the manufacturer’s descriptions using the HotStarTaq PLUS DNA Polymerase kit (Qiagen) on a PTC-200 Peltier Thermal Cycler (MS Research). To catch cDNA amplification in the exponential phase, experiments were run for 30 cycles only as follows: initial heat activation at 95 °C for 5 min, denaturation at 94 °C for 30 sec, annealing at primer-specific temperatures listed in table below for 40 sec, extension at 72 °C for 1 min, final extension at 72 °C for 7 min. qRT-PCR was performed using the SYBR Green method according to the manufacturers’ descriptions (Roche), using the default SYBR Green program on a Roche LightCycler 480 (Roche). All experiments were performed on at least three independent biological replicates and each sample was run in triplicate for qRT-PCR. Data were normalized to housekeeping genes for comparisons and fold change calculated by the 2ΔΔCT method. Supplementary Figure 2A shows technical replicates. As ATOH1 was not detected by the end of the qRT-PCR at cycle number 45 (but the house keeping gene was) in the EGFP-samples, 45 was assigned as the ATOH1 cycle number value (CT) to signify a non-detected signal and calculate the fold change in the EGFP+ samples compared to EGFP-samples. For primer sequences and annealing temperatures see table below. Primers were designed using Primer3 (https://bioinfo.ut.ee/primer3-0.4.0/) and validated to give single amplicons in a concentration dependent manner at temperatures used.

**Methods Table 1:**
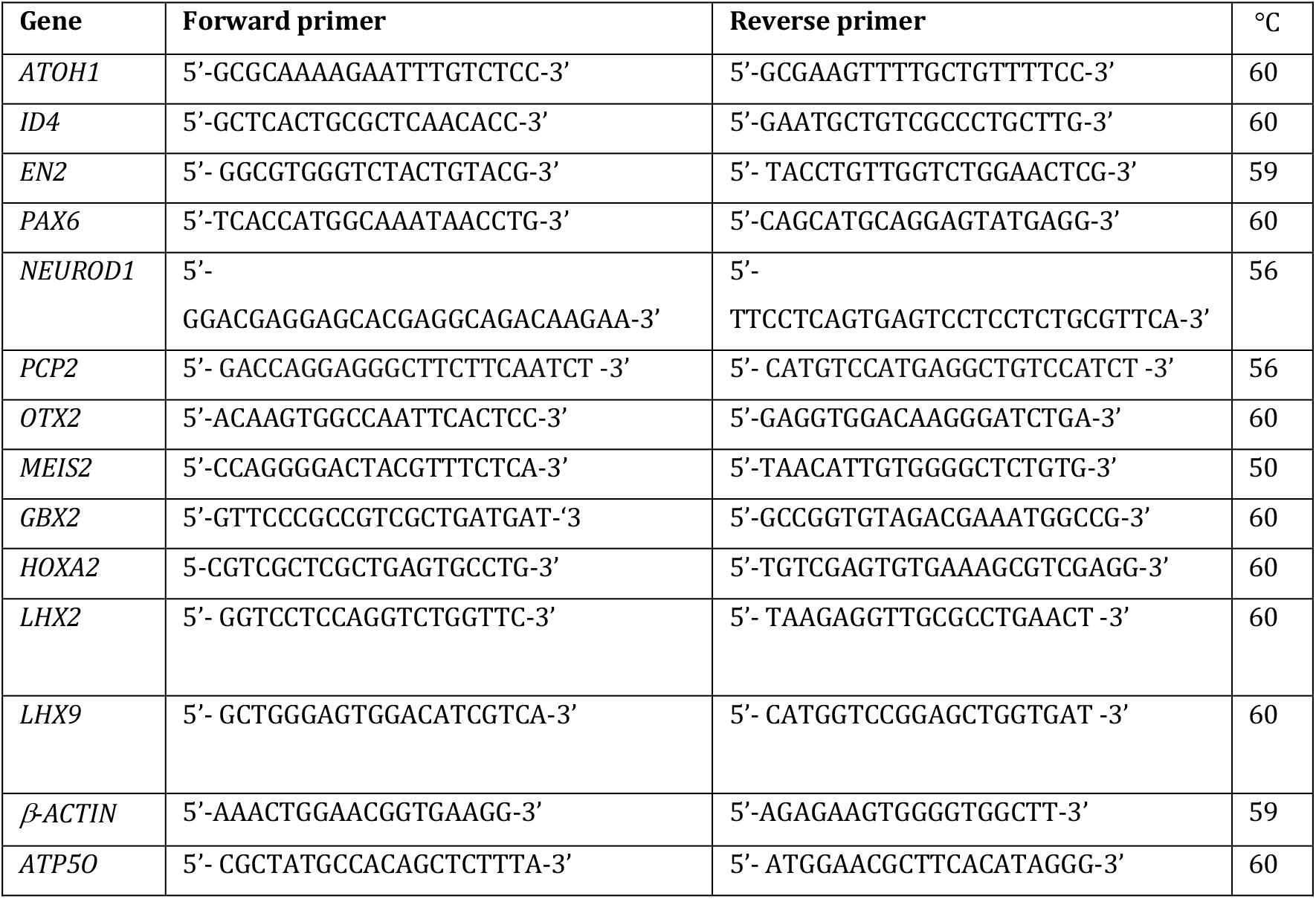
Table of primers

### Immunocyto/histochemistry

Cells grown on 1.5 thickness cover glass (Fisher) were fixed for 15 min at RT with 4% PFA and washed 3x PBS. Cells were then blocked in PBS containing 1% normal horse serum (Gibco) and 0.1% Triton for 1 hr and then incubated with primary antibodies in blocking solution for 1hr at RT to overnight at 4°C. Cells were washed 3x PBS for 10 min and Alexa Fluor® conjugated secondary antibody incubations were performed in blocking solution for 1 hr at RT. Dapi was sometimes added as nuclear counterstain (1 μg/ml, Molecular Probes). For immunohistochemistry on frozen brain sections, human cerebella were fixed and embedded as described earlier. Mouse brains at E15.5 and P0 were fixed in 4% PFA overnight at 4°C, then cryoprotected in 20% sucrose overnight at 4°C and embedded in OCT. 14 μm thick sagittal sections were prepared for all brains on a Leica CM 3050S cryostat. Frozen sections were thawed, postfixed for 10 min in 4% PFA at RT and immunohistochemistry was carried out as above, except that the blocking solution contained 10% normal horse serum (Gibco) and 0.2% Triton in PBS. For analysis of transplantation experiments, brains were fixed in 4% PFA overnight at 4°C, washed in PBS, and embedded in 3% agarose. 50 μm thick vibratome sections were postfixed with 4% PFA for 15 min at RT followed by blocking in 10% normal horse serum (Gibco) and 0.2% Triton in PBS overnight at 4°C. Primary antibody incubations were carried out in blocking solution for two nights at 4°C followed by extensive washes (4x 15 min each) in PBS containing 0.1% Triton and sections were then incubated in secondary antibodies overnight at 4°C. Sections/cells were mounted with ProLong® Gold anti-fade mounting media (Invitrogen) and 1.5 thickness Fisherbrand cover glass. For antibody sources and dilutions see table below.

**Methods Table 2:** Table of antibodies

| Antibody | Species | Source | Dilution | Catalogue # |
| --- | --- | --- | --- | --- |
| Calretinin | Rabbit | Swant | 1:1000 | 7699/4 |
| EN2 (C19) | Goat | Santa Cruz | 1:50 | SC-8111 |
| GFAP | Chicken | Encor Biotechnology | 1:1500 | CPCA-GFAP |
| GFP | Rabbit | Invitrogen | 1:500 | A-111122 |
| GFP | Chicken | Aves labs | 1:1000 | GFP-1020 |
| HuNu (anti human nuclei) | Mouse | Millipore | 1:100 | MAB1281 |
| Ki67 | Rabbit | Vector laboratories | 1:100 | VP-RM04 |
| Ki67 | Rabbit | EnCor | 1:1000 | RPCA-Ki67 |
| MAP2 | Chicken | Abcam | 1:1000 | ab5392 |
| NEUN | Mouse | Millipore | 1:100 | MAB377 |
| NEUROD1 | Mouse | BD Pharmingen | 1:300 | 563000 |
| PAX6 | Rabbit | Biolegend | 1:300 | 901301 |
| TAG1 | Mouse IgM | T. Jessell | 1:2 | N/A |
| Synaptophysin | Mouse IgM | Millipore | 1:500 | MAB329 |
| Anti goat Alexa Fluor® 633 | Donkey | Invitrogen | 1:300 | A21082 |
| Anti rabbit Alexa Fluor® 555 | Donkey | Invitrogen | 1:300 | A-31572 |
| Anti mouse IgM Alexa Fluor® 488 | Goat | Invitrogen | 1:300 | A-21042 |
| Anti mouse Alexa Fluor® 555 | Donkey | Invitrogen | 1:300 | A-31570 |
| Anti chicken IgY 488 | Donkey | Jackson Immunoresearch | 1:300 | 703-545-155 |
| Anti chicken IgY Cy3 |  | Jackson Immunoresearch | 1:300 | 703-165-155 |
| Anti rabbit Alexa Fluor® 647 | Donkey | Invitrogen | 1:300 | A-31573 |
| Anti mouse 405 | Donkey | Abcam | 1:300 | Ab175658 |
| Anti mouse Alexa Fluor® 647 | Donkey | Invitrogen | 1:300 | A-31571 |
| Anti mouse Alexa Fluor® 488 | Donkey | Invitrogen | 1:300 | A-21202 |

### Proliferation assay

Cell proliferation was measured by EdU incorporation using the Click-iT™ EdU Cell Proliferation Kit for Imaging (Invitrogen, C10338) according to the manufacturer’s description. On day 11 of differentiation, cells from the *ATOH1-EGFP* line were gently scraped off Transwell membranes and plated on poly-D-lysine (Millipore, A-003-E) and Laminin (Invitrogen, 23017-015)-coated glass coverslips and differentiation was resumed as described previously (Fig. 1D). On day 28, EdU was added to the culture medium according to the manufacturer’s description and cells were treated with either SAG (0.5μM, Cayman Chemicals 11914), or DMSO. Cells were fixed with 4% PFA (15 min at RT) on day 30 and processed for Click-iT EdU detection and immunocytochemistry. See below for quantification and statistical analysis.

### Transplantation of hPSC-derived cerebellar granule cells in the mouse cerebellum

All procedures were approved by the Rockefeller University Institutional Animal Care and Use Committee. TAG1+ cells were isolated by MACS on DIV 28-32 as described earlier. Cells were counted and resuspended in “transplantation medium” containing BME, Gibco; 10% v/v horse serum, Invitrogen 26050-088; 0.9% v/v glucose, Sigma; and 0.5% w/v Fast Green for visualization of injection volumes. Cells were kept on ice while mouse pups were prepared for injections (up to 2 hrs). Neonatal mouse pups (P1-4) were cryo-anesthetized, the heads were cleaned with alcohol and a 1μl single cell suspension (5 x 10^6^ cells/ml) was manually injected directly into the left cerebellar hemisphere using a glass microcapillary (Eppendorf, 5195 000.079) controlled with an Eppendorf CellTram Vario manual microinjector. The procedure was performed under a Zeiss OMPI-1FC surgical microscope (ENT) with Zeiss eyepieces. The capillary directly pierced through both skin and skull. The positioning of the capillary was guided by the left earlobe and Lambda and the tip of the capillary was placed just under the skull to target the cerebellar surface. The capillary was held in place for 1 min after the completion of injection to minimize backflow after which it was gently pulled out and the pups was warmed up on a heating pad (Sunbeam) before being returned to their mothers. Injected animals were analyzed 48hrs post injection. Brains were dissected out and fixed overnight in 4% PFA. The entire cerebellum of each animal (N=5 analyzed) was sectioned coronally at 50μm thickness on a vibratome (Leica VT 1000S) and all sections were processed for immunohistochemistry using a human nuclear antigen antibody (HuNu) to detect human cells plus additional antibodies as described in manuscript.

### TRAP and RNA sequencing

TRAP was performed as previously reported (Heiman et al., 2014) on three independent differentiation experiments at DIV28. Briefly, polysomes were stabilized by adding 100 μg/ml cycloheximide to cell culture media for 10 minutes prior to homogenization of cells with polysome extraction buffer. Following clearing by centrifugation, supernatants were incubated at 4°C with end-over-end rotation for 16-18 hrs with biotinylated Streptavidin T1 Dynabeads (ThermoFisher, 65601) previously conjugated with GFP antibodies (Sloan Kettering Institute Antibody Core, HtzGFP-19C8 and HtzGFP-19F7). The beads were collected on a magnetic rack, washed, and resuspended in lysis buffer with β-mercaptoethanol (Agilent, 400753) to extract bound RNA from polysomes. RNA was purified using the RNeasy micro kit (Qiagen, 74004). RNA quantity and quality were measured using an Agilent 2100 Bioanalyzer with the 6000 Pico Kit (Agilent, 5067-1513). Full-length cDNA was prepared using Clontech’s SMART-Seq v4 Ultra Low Input RNA Kit (634888) from 0.5 ng RNA with an RIN≥8.3. 1 ng cDNA was then used to prepare libraries using Illumina Nextera XT DNA sample preparation kit (FC-131-1024). Libraries with unique barcodes were pooled at equal molar ratios and sequenced on Illumina NextSeq 500 sequencer to generate 75 bp single reads, following the manufacturer’s protocol.

### RNA sequencing analysis

Sequence and transcript coordinates for human hg19 UCSC genome and gene models were retrieved from the Bioconductor Bsgenome.Hsapiens.UCSC.hg19 (version 1.4.0) and TxDb.Hsapiens.UCSC.hg19.knownGene (version 3.2.2) Bioconductor libraries respectively. Transcript expressions were calculated using the Salmon quantification software (Patro et al., 2017) (version 0.8.2) from raw FastQ files. Gene expression levels as TPMs and counts were retrieved using Tximport (Love et al., 2016) (version 1.8.0). Normalization and rlog transformation of raw read counts in genes were performed using DESeq2 (Love et al., 2018) (version 1.20.0). For visualization in genome browsers, RNA-seq reads were aligned to the genome using Rsubread’s subjunc method (version 1.30.6) (Liao et al., 2013) and exported as bigWigs normalized to reads per million using the rtracklayer package (version 1.40.6). Genes significantly enriched and depleted in IP over input were identified using DESeq2 at padj cutoff of 0.05 and absolute log fold change cut-offs of both 0 and 2. The PsychEncode’s ‘Human mRNA-seq processed data’ as counts was retrieved from the PsychEncode’s portal (http://development.psychencode.org). Analysis of significantly enriched or depleted gene sets (absolute logFC > 2, padj < 0.05) were performed using GSVA (version 1.34.0) (Hanzelmann et al., 2013) against DESeq2 normalised PsychEncode midbrain and cerebellum RNA-seq counts. Visualization of genes and gene-sets as heat maps was performed using the Pheatmap R package (version 1.0.10) (Subramanian et al., 2005). GO term enrichment was obtained for all genes enriched in IP above input (logFC > 0, padj < 0.05) using the fisher test in the topGO Bioconductor package and ranked using the elim algorithm and functional annotation from the org.Hs.eg.db Bioconductor package (version 3.10).

### Imaging

Single z-plane images (512 x 512 pixels) were acquired using an inverted Zeiss LSM 880 NLO laser scanning confocal microscope with a Plan-Apochromat 40x/1.4 NA objective oil immersion lens fitted with a Nomarski prism, and HeNe and Argon lasers for excitation at 405, 488, 561, and 633 nm. Phase contrast images in Supp Fig. 1D were acquired with a 20X objective on a Leica DMIL LED microscope fitted with a Leica MC120 HD camera.

### Quantification and statistical analyses

For immunocyto/histochemistry experiments, cells were manually counted in ImageJ (version 2.1.0/1.53c) on single z-plane confocal images (512 x 512 pixels) from at least three independent experiments. The percentages of PAX6, EN2, and EGFP positive cells were calculated per PAX6, EN2, and EGFP populations as indicated in Fig. 1F. Multiple representative images per coverslip were quantified and a total of 1787 EN2^+^, 3487 GFP^+^, and 2845 PAX6^+^ cells were counted. The percentage of EdU/EGFP double positive cells were calculated as a subset of the Dapi population. A total of 84 images (6209 EdU^+^ cells) were analyzed with five outliers removed. Data were log transformed for normality and the SAG and control treatment groups were compared by ANCOVA with the number of Dapi^+^ cells as a covariate. The number of TAG1^+^ cells were counted after MACS using a haemocytometer under a Leica microscope (Leica DMIL LED) and expressed as a percentage of the cells at the start of the sort (input). The percentage of NEUROD1 positive cells at the pial surface of the cerebellum were counted and normalized to the number of Dapi^+^ nuclei along the length of the pial surface. Multiple sagittal sections from 2 brains per time point and species were analyzed with the total numbers of cells/sample indicated in the description of Supplementary Fig. 4.

## Data availability

RNAseq data can be found at https://www.ncbi.nlm.nih.gov/geo/ submission number: GSE163710. Dataset 1 contains DESeq2 analysis of *ATOH1-EGFP-L10a* TRAP IP versus input.

## Acknowledgements

We thank Dr. Ali Brivanlou and Dr. Zeeshan Ozair (The Rockefeller University) for the provision of vital reagents and critical discussions throughout the project, Dr. Nathaniel Heintz and Jie Xing (The Rockefeller University) for the kind gift of the EGFP-L10a containing construct, helpful discussions regarding the adaptation of bacTRAP methodology for use in hPSCs and technical support. We also thank Dr. Jane Johnson (UT Southwestern) for the kind gift of the J2XnGFP plasmid and the late Dr. Tom Jessell (Columbia University) for the provision of the TAG1 antibody. We are also grateful to staff at the Rockefeller Core facilities including imaging, flow cytometry, and high-throughput sequencing. Finally, we thank members of the Hatten lab for helpful discussions. This work was supported by NIH 1R21NS093540-01 (M.E.H), a Rockefeller University Center for Clinical and Translational Science Pilot grant (M.E.H, H.B), a Tri-Institutional Stem Cell Initiative grant from the Starr Foundation (M.E.H), and a gift from the Renate Hans and Maria Hofmann Trust (M.E.H).

