## Supplemental Table 1 for "Altered temporal sequence of transcriptional regulators in the generation of human cerebellar granule cells"

| GO ID | GO term | Annotated | Significant | Expected | Rank in resul | Classic | Elim | Weight | |
|---|---|---|---|---|---|---|---|---|---|
| GO:0006613 | cotranslational protein targeting to membrane | 96 | 87 | 28.02 | 1 | < 1e-30 | < 1e-30 | 0.02662 | 7.50E-14 |
| GO:0000184 | nuclear-transcribed mRNA catabolic proce... | 119 | 98 | 34.74 | 2 | < 1e-30 | < 1e-30 | < 1e-30 | 2.20E-07 |
| GO:0019080 | viral gene expression | 186 | 123 | 54.3 | 3 | 1.20E-25 | 4.80E-28 | 0.01712 | 6.40E-14 |
| GO:0019058 | viral life cycle | 447 | 228 | 130.49 | 4 | 8.70E-23 | 9.90E-24 | 6.80E-25 | 3.20E-08 |
| GO:0044033 | multi-organism metabolic process | 212 | 131 | 61.89 | 5 | 4.10E-23 | 4.10E-23 | 4.10E-23 | 1.50E-20 |
| GO:0006412 | translation | 619 | 308 | 180.7 | 6 | 4.70E-28 | 1.10E-21 | 0.04048 | 2.10E-20 |
| GO:0006364 | rRNA processing | 253 | 148 | 73.86 | 7 | 1.50E-22 | 1.40E-21 | 4.60E-15 | 8.30E-07 |
| GO:0006119 | oxidative phosphorylation | 118 | 81 | 34.45 | 8 | 7.20E-19 | 7.20E-19 | 0.01614 | 1.80E-14 |
| GO:0044764 | multi-organism cellular process | 952 | 395 | 277.91 | 9 | 5.00E-17 | 5.80E-17 | 5.00E-17 | 8.60E-16 |
| GO:0033108 | mitochondrial respiratory chain complex ... | 83 | 57 | 24.23 | 10 | 1.10E-13 | 1.10E-13 | 0.00102 | 1.60E-08 |
| GO:0000398 | mRNA splicing, via spliceosome | 309 | 145 | 90.2 | 11 | 2.30E-11 | 2.70E-10 | 6.90E-07 | 0.00052 |
| GO:0007411 | axon guidance | 221 | 112 | 64.51 | 12 | 1.20E-11 | 7.50E-10 | 2.40E-08 | 0.00088 |
| GO:0051649 | establishment of localization in cell | 2046 | 845 | 597.26 | 13 | < 1e-30 | 8.00E-10 | 0.03914 | 6.50E-21 |
| GO:0030198 | extracellular matrix organization | 313 | 148 | 91.37 | 14 | 7.10E-12 | 3.00E-09 | 2.50E-08 | 0.0027 |
| GO:0051961 | negative regulation of nervous system de... | 256 | 124 | 74.73 | 15 | 5.00E-11 | 7.20E-09 | 0.0078 | 4.20E-05 |
| GO:1903047 | mitotic cell cycle process | 900 | 367 | 262.73 | 16 | 1.50E-14 | 1.30E-08 | 0.00016 | 0.00024 |
| GO:0071363 | cellular response to growth factor stimu... | 617 | 264 | 180.11 | 17 | 1.70E-13 | 1.30E-08 | 0.00278 | 1.50E-08 |
| GO:0009952 | anterior/posterior pattern specification | 202 | 99 | 58.97 | 18 | 2.00E-09 | 1.60E-08 | 2.00E-09 | 0.00896 |
| GO:0007417 | central nervous system development | 895 | 363 | 261.27 | 19 | 5.20E-14 | 2.00E-08 | 0.00011 | 1.60E-05 |
| GO:0072594 | establishment of protein localization to... | 657 | 311 | 191.79 | 20 | 1.00E-23 | 2.70E-08 | 0.00302 | 1.00E-08 |
| GO:0006914 | autophagy | 448 | 193 | 130.78 | 21 | 1.70E-10 | 2.70E-08 | 0.049 | 0.01591 |
| GO:0048706 | embryonic skeletal system development | 122 | 67 | 35.61 | 22 | 2.30E-09 | 6.50E-08 | 0.02204 | 0.00069 |
| GO:0042274 | ribosomal small subunit biogenesis | 64 | 39 | 18.68 | 23 | 1.30E-07 | 1.30E-07 | 0.23874 | 0.07409 |
| GO:0016055 | Wnt signaling pathway | 469 | 192 | 136.91 | 24 | 2.40E-08 | 1.70E-07 | 0.02202 | 0.0083 |
| GO:0048705 | skeletal system morphogenesis | 209 | 103 | 61.01 | 25 | 6.30E-10 | 3.40E-07 | 0.06834 | 0.00224 |
| GO:0048193 | Golgi vesicle transport | 331 | 151 | 96.62 | 26 | 1.20E-10 | 3.80E-07 | 0.00972 | 4.40E-05 |
| GO:0043066 | negative regulation of apoptotic process | 808 | 311 | 235.87 | 27 | 3.70E-09 | 5.40E-07 | 0.00066 | 0.03687 |
| GO:0042273 | ribosomal large subunit biogenesis | 60 | 36 | 17.52 | 28 | 6.60E-07 | 6.60E-07 | 0.03329 | 0.11008 |
| GO:0043161 | proteasome-mediated ubiquitin-dependent ... | 376 | 155 | 109.76 | 29 | 3.20E-07 | 8.80E-07 | 0.39402 | 0.01383 |
| GO:0051301 | cell division | 563 | 229 | 164.35 | 30 | 2.10E-09 | 1.20E-06 | 1.80E-07 | 1.50E-08 |
| GO:0001764 | neuron migration | 136 | 66 | 39.7 | 31 | 1.40E-06 | 1.50E-06 | 3.80E-05 | 0.00472 |
| GO:0010628 | positive regulation of gene expression | 1723 | 596 | 502.98 | 32 | 1.60E-07 | 1.70E-06 | 0.00016 | 0.24635 |
| GO:0032392 | DNA geometric change | 84 | 45 | 24.52 | 33 | 2.40E-06 | 2.40E-06 | 0.13488 | 0.00032 |
| GO:0036293 | response to decreased oxygen levels | 340 | 140 | 99.25 | 34 | 1.30E-06 | 2.40E-06 | 0.17545 | 0.37417 |
| GO:0060322 | head development | 705 | 286 | 205.8 | 35 | 2.80E-11 | 2.80E-06 | 0.03897 | 3.20E-06 |
| GO:0007041 | lysosomal transport | 82 | 44 | 23.94 | 36 | 2.90E-06 | 2.90E-06 | 0.00365 | 0.62205 |
| GO:0001503 | ossification | 372 | 162 | 108.59 | 37 | 1.90E-09 | 3.00E-06 | 0.0003 | 2.10E-09 |
| GO:0061548 | ganglion development | 13 | 12 | 3.79 | 38 | 3.60E-06 | 3.60E-06 | 0.49797 | 2.40E-05 |
| GO:0051170 | import into nucleus | 291 | 123 | 84.95 | 39 | 1.10E-06 | 3.70E-06 | 0.00172 | 0.29048 |
| GO:0014812 | muscle cell migration | 74 | 40 | 21.6 | 40 | 6.40E-06 | 4.00E-06 | 0.00367 | 0.00092 |
| GO:2000058 | regulation of ubiquitin-dependent protei... | 207 | 91 | 60.43 | 41 | 4.00E-06 | 7.30E-06 | 0.08417 | 0.01815 |
| GO:0000291 | nuclear-transcribed mRNA catabolic proce... | 35 | 23 | 10.22 | 42 | 8.10E-06 | 8.10E-06 | 0.49757 | 0.70948 |
| GO:0007169 | transmembrane receptor protein tyrosine ... | 680 | 259 | 198.5 | 43 | 2.20E-07 | 8.40E-06 | 0.00012 | 0.68296 |
| GO:0048666 | neuron development | 973 | 393 | 284.04 | 44 | 9.00E-15 | 1.10E-05 | 0.00663 | 0.00012 |
| GO:0033554 | cellular response to stress | 1801 | 689 | 525.74 | 45 | 1.20E-18 | 1.40E-05 | 0.00062 | 9.50E-22 |
| GO:0002062 | chondrocyte differentiation | 96 | 49 | 28.02 | 46 | 5.40E-06 | 1.40E-05 | 0.00037 | 9.00E-05 |
| GO:0043065 | positive regulation of apoptotic process | 572 | 221 | 166.98 | 47 | 5.10E-07 | 1.60E-05 | 3.20E-05 | 0.07929 |
| GO:0043583 | ear development | 204 | 89 | 59.55 | 48 | 7.30E-06 | 1.80E-05 | 0.00237 | 0.06719 |
| GO:0070849 | response to epidermal growth factor | 43 | 26 | 12.55 | 49 | 1.90E-05 | 1.90E-05 | 0.00719 | 0.00054 |
| GO:1903321 | negative regulation of protein modificat... | 144 | 66 | 42.04 | 50 | 1.60E-05 | 2.40E-05 | 0.01415 | 0.00131 |
